# Protein tyrosine phosphatase-PEST (PTP-PEST) mediates hypoxia-induced endothelial autophagy and angiogenesis through AMPK activation

**DOI:** 10.1101/2020.06.15.152942

**Authors:** Shivam Chandel, Amrutha Manikandan, Nikunj Mehta, Abel Arul Nathan, Rakesh Kumar Tiwari, Samar Bhallabha Mohapatra, Mahesh Chandran, Abdul Jaleel, Narayanan Manoj, Madhulika Dixit

## Abstract

Global and endothelial loss of PTP-PEST is associated with impaired cardiovascular development and embryonic lethality. Although hypoxia is implicated in vascular morphogenesis and remodelling, its effect on PTP-PEST remains unexplored. Here we report that hypoxia (1% oxygen)increases protein levels and catalytic activity of PTP-PEST in primary endothelial cells. Immunoprecipitation followed by mass spectrometry (LC/MS/MS) revealed that AMP-activated protein kinase alpha subunits (AMPK α_1_ and α_2_) interact with PTP-PEST under normoxia but not in hypoxia. Co-immunoprecipitation experiments confirmed this observation and determined that AMPK α subunits interact with the catalytic domain of PTP-PEST. Knock-down of PTP-PEST abrogated hypoxia mediated tyrosine dephosphorylation and activation of AMPK (Thr^172^ phosphorylation). Absence of PTP-PEST also blocked hypoxia-induced autophagy (measured as LC3 degradation and puncta formation) which was rescued by AMPK activator, metformin (500µM). Since endothelial autophagy is a pre-requisite for angiogenesis, knock-down of PTP-PEST also attenuated endothelial cell migration and capillary tube formation with autophagy inducer rapamycin (200nM) rescuing these effects. In conclusion, this work identifies for the first time PTP-PEST as a regulator of hypoxia-induced AMPK activation and endothelial autophagy to promote angiogenesis.

## Introduction

Physiological hypoxia is a potent agonist for embryonic development and post-natal angiogenesis. Oxygen concentrations ranging from 1% to 5% are observed in the uterine environment between embryonic days 3.5 to 14.5 (E3.5-E14.5) to facilitate development of placenta. Similarly, in response to hypoxia (<2% oxygen), endocardial and vascular endothelial cells mediate formation of foetal heart and vasculature respectively in mice between embryonic days E7.5 to E15 [1;2]. Post-natal angiogenesis as seen in female reproductive tract during menstrual cycle in humans, as well as formation of collateral circulation to over-come coronary artery blocks also depend on hypoxia-induced endothelial signalling. Other than the known classical mediators of angiogenesis, multiple studies in rodents have reported increased activity of cytosolic protein tyrosine phosphatases (PTPs) at the sites of post-natal angiogenesis, including ischemic myocardium and skeletal muscles [3;4]. An increase in the cytosolic PTP activity during hypoxia is also seen in the cerebral cortex of new born piglets [5]. Paradoxically, others have shown that non-selective PTP inhibitor, sodium orthovanadate, enhances VEGFR2 signalling and capillary morphogenesis [3;6]. Although these studies allude to the involvement of different PTPs in hypoxia-induced angiogenic signalling, barring the involvement of few, for instance VE-PTP (a receptor PTP), the identity of hypoxia responsive cytosolic angiogenic PTPs remains largely unknown.

Human genome encodes 38 classical PTPs which are specific for tyrosine residues [7]. Among these, PTP-PEST is a ubiquitously expressed cytosolic PTP with an N-terminal catalytic domain (1-300 amino acids) and a regulatory C-terminal PEST domain (301-780 amino acids) containing four PEST motifs. The latter plays a crucial role in protein-protein interaction allowing the enzyme to interact with its known substrates such as Cas, Paxillin, FAK and Pyk2 in addition to being a protein degradation signal [8-10]. The catalytic domain harbours the conserved phosphatase ‘HC(X)_5_R’ motif surrounded by the ‘WPD loop’ and the ‘Q loop’ on one side to assist in catalysis and by the ‘P-Tyr-loop’on the other side to regulate substrate recognition and specificity [11]. Both global and endothelial deficiency of PTP-PEST in mice leads to embryonic lethality between E9.5-E10.5 days due to defective heart formation and impaired endothelial network in the yolk sac [12;13]. It is worth noting that this stage in embryonic development coincides with hypoxia-induced morphogenesis [1;2]. PTP-PEST appears to be crucial for vascular development even in humans since a partial deletion of PTP-PEST is associated with disrupted aortic arch development [14]. Intriguingly, in a recent study, enhanced endothelial expression of PTP-PST was indeed observed in the vascularized core of glioblastoma tumours with others demonstrating involvement of PTP-PEST in integrin mediated endothelial cell adhesion and migration [13;15]. Based on these observations which illustrate an essential role of PTP-PEST in vascular development and given the fact that hypoxia is indispensable for cardiovascular development and angiogenesis, in the present study we set out to determine the functional role of PTP-PEST in hypoxia-induced endothelial responses. We assessed the effect of hypoxia on the expression, activity and binding partners of PTP-PEST in primary human umbilical vein derived endothelial cells (HUVECs).

## Materials and methods

Experimental procedure involving isolation of endothelial cells from umbilical cords was approved by the IIT-Madras Institutional Ethics Committeeas per the Indian Council of Medical Research (ICMR), Government of India guidelines. These guidelines are in accordance with the declaration of Helsinki, which was revised in 2000.

### Antibodies and Chemicals

Tissue culture grade plastic ware was from Tarsons Products Pvt. Ltd., Kolkata, India. Antibodies against HIF-1α, phospho-AMPK, total AMPK, LC3, phospho-ULK1, total ULK1, phospho-ACC, total ACC, PTP-PEST (AG10) and anti-phospho-tyrosine (P-Tyr-102) antibody were obtained from Cell Signaling Technology (CST), Boston, MA, USA. Dulbecco’s modified Eagle’s medium (DMEM) and MCDB131 were purchased from HiMedia™, Mumbai, India and fetal bovine serum (FBS) was from Gibco, Thermo Fisher Scientific, Waltham, MA, USA. Unless specified otherwise rest of the molecular biology and biochemistry reagents were from Sigma, St. Louis, MO, USA.

### Cell culture

HUVECs were isolated from freshly collected umbilical cords by collagenase digestion. They were cultured in fibronectin coated T25 flasks until monolayer was formed. MCDB131 medium with endothelial growth supplements was used for culturing HUVECs. All the experiments were performed in passage one. For cell lines HEK293, HeLa, Huh7 and HASMC, cells were cultured in DMEM with 10% FBS. For hypoxia exposure, cells were incubated in hypoxia incubator (5% CO_2_, 1% O_2_ and 95% relative humidity) for specified duration. After degassing, the cell culture media used for hypoxia experiments was pre-equilibrated to the hypoxic environment prior to start of the experiment.

### Immunoblotting

All the buffers used for washing and lysing of cells were degassed before use. Cells were washed with 1X cold PBS and lysed in buffer containing 50mM Tris-HCl (pH 7.5), 150mM NaCl, 1mM EDTA, 1mM EGTA, 1% TritonX-100, 0.1% SDS, 1% sodium deoxycholate, 1mM PMSF and protease inhibitor cocktail from Sigma (St. Louis, MO, USA) followed by sonication for 1 min (30% amplitude, 3 second on/off cycle). 50 µg worth total protein was resolved through 8% SDS-PAGE and transferred onto PVDF membrane. Blotted proteins were incubated with corresponding primary antibodies at 4°C overnight followed by incubation with horseradish peroxidase conjugated secondary antibodies (Jackson ImmunoResearch, West Grove, PA, USA). Amersham ECL Western blotting detection reagent (GE Healthcare Life Sciences, USA) was used for detection. Image J version 1.45 (NIH, Bethesda, MD, USA) was used for densitometry analysis.

### Immunoprecipitation and phosphatase assay

Cells were lysed in ice cold buffer containing 50mM Tris-HCl (pH 7.5), 150mM NaCl, 1mM EDTA, 1mM EGTA, 1% Triton-X 100, 2.5mM sodium pyrophosphate, 1mM β-glycerophosphate, 1mM sodium orthovanadate, protease inhibitor cocktail (Sigma, St. Louis, MO, USA) and phosphatase inhibitor cocktail 2 (Sigma, St. Louis, MO, USA) followed by 30 seconds of sonication. 300 µg worth total protein was pre-cleared using protein A/G sepharose beads.This was followed by overnight incubation with PTP-PEST AG10 monoclonal antibody at 4°C. Protein-antibody complex was pulled down using protein A/G sepharose beads in sample buffer.Specific pull down of PTP-PEST was confirmed by employing corresponding IgG isotype antibody as negative control. For phosphatase assay elution was performed in assay buffer. Phosphatase assay was carried out in Na-acetate buffer [2mM EDTAand 1mM DTT, pH 5.0] at 30°C for 2 hours and 50mM pNPP was used as substrate. The catalytic activity was determined spectrophotometrically by measuring the amount of pNP (para-nitrophenol) generated from pNPP. pNP released was quantified by measuring absorbance at 420 nm.

### Lentivirus production and transduction

pcDNA3.1 Flag-PTP-PEST plasmid construct was a kind gift from Dr.Zhimin Lu, Department of Neuro-oncology, The University of Texas M. D. Anderson Cancer Center, Houston, USA [16]. PTP-PEST was sub-cloned in pENTR4-GST-6P1 (Addgene#17741) lentivirus entry vector backbone at BamH1 and XbaI restriction sites followed by LR clonase mediated gateway recombination in 3^rd^ generation lentivirus destination vector pLenti CMV Puro Dest (Addgene#17452). shRNA oligos(Oligo 1:5’-GATCCCACCAGAAGAATCCCAGAATCTCGAGATTCTGGGATTCTTCTGGTGGTTTTTG-3’ and Oligo 2:5’-AGCTCAAAAACCACCAGAAGAATCCCAGAATCTCGAGATTCTGGGATTCTTCTGGTGG-3’) were annealed in annealing buffer (10mM Tris pH 8.0, 50mM NaCl, 1mM EDTA) and cloned in pENTR/pSUPER^+^ entry vector (Addgene#17338) which was recombined into pLenti X1 Puro DEST (Addgene#17297). These lentivirus entry and destination plasmids were a gift from Eric Campeau and Paul Kaufman [17]. For lentivirus production, recombined expression vectors were co-transfected with packaging plasmids pLP1, pLP2 and pVSVG in 293T cells. Supernatant was collected 48 hours and 72 hours post-transfection and was concentrated through ultracentrifugation. Lentiviral transduction was performed in HUVECs (P_0_) in low serum (5%) endothelial growth medium at 70% confluency in presence of polybrene (6µg/ml) for 8 hours. This was followed by a second round of transduction. 48 hours post-transduction, puromycin (2µg/ml) selection was performed for next 3 days followed by splitting of cells for experimental treatment.

### Immunofluorescence imaging

Following appropriate experimental treatments, HUVECs were washed with 1X PBS, followed by paraformaldehyde (0.4%) fixing and cells were made permeable with 0.25% Triton X. Cells were blocked with 5% serum in 1X PBS followed by overnight staining with primary antibodies at 4°C. Cells were then washed and incubated with fluorophore conjugated secondary antibodies (Thermo Fisher Scientific, Waltham, MA, USA). Nuclei were counterstained with DAPI and images were captured in LSM 700 Zeiss confocal microscope.

### Mass spectrometry

HUVECs were transduced with N-terminal GST tagged PTP-PEST lentiviral particles. Post-transduction, same pool of cells was split and exposed to either normoxia or hypoxia for 24 hours. GST tagged PTP-PEST was pulled down via immunoprecipitation using GST antibody (Cell Signaling Technology, USA). Immunoprecipitated proteins were loaded on to SDS-PAGE. SDS-PAGE was done in such a way that the electrophoresis run was stopped the moment the entire sample entered the resolving gel. Each band approximately 5mm in size, visible after Coommassie Blue staining was excised and used for proteomics analysis. The proteomic profiling was performed by liquid chromatography mass spectrometry (LC/MS/MS) at the Mass Spectrometry and Proteomics Core facility of RGCB, Thiruvananthpuram. Briefly, the excised gel pieces were subjected to in-gel trypsin digestion using sequence grade trypsin (Sigma) as per Shevchenko et al 2006 [18]. The LC/MS/MS analyses of the extracted tryptic peptides were performed in a SYNAPT G2 High Definition Mass Spectrometer (Waters, Manchester, UK), which is connected to a nanoACQUITY UPLC® chromatographic system (Waters) for the separation of peptides. The LC/MS acquired raw data was analyzed by Progenesis QI for Proteomics V3.0 (NonLinear Dynamics, Waters) for protein identification using the Human protein database downloaded from UniProt.

### Wound healing assay

HUVECs were cultured as tight monolayer and a scratch was made to create a wound. Immediately after wound creation, scrapped cells were washed off and images were taken at the start as 0 hour. HUVECs were then incubated in presence of 5mM hydroxyurea (anti-proliferative molecule) for next 24 hours either in normoxia or 1% hypoxia condition and images were retaken at the same locations following incubation to assess migration. Area of wound closure in 24 hours was calculated by employing ImageJ software (NIH, Bethesda, MD, USA).

### *In vitro* tube formation assay

Wells in a 96 well plate were coated with 80 µl of growth factor free Matrigel and allowed to polymerize in CO_2_ cell culture incubator. Following appropriate experimental treatment (hypoxia and/or knock-down) 15 × 10^3^ cells were seeded onto each well and incubated for 16 hours in CO_2_ cell culture incubator followed by imaging. Tube networks were analyzed by using angioanalyser feature of ImageJ software (NIH, Bethesda, MD, USA).

### Site directed mutagenesis

Wild type PTP-PEST containing first 300 amino acids was PCR amplified using primers (Forward primer: 5’-CGCGGATCCATGGAGCAAGTGGAGATC-3’, Reverse primer: 5’-CCGCTCGAGTCATAGTTGTAGCTGTTTTTC-3’) and cloned in pET28a(+) bacterial expression vector between BamH1 and Xho1 restriction sites. Site directed mutagenesis was carried out to generate C231S mutant using following primers, forward primer: 5’-TGTATTCATTCCAGTGCAGGCTG-3’, reverse primer: 5’-CAGCCTGCACTGGAATGAATACA-3’. Polymerase chain reaction (PCR) was performed using 10ng of His tagged PTP-PEST (WT) plasmid as template and followed by Dpn1 (New England Biolabs, UK) digestion for 2 hours at 37°C. Dpn1 digested PCR product was transformed into *E. coli* DH5α ultra-competent cells. Positive clones were selected by kanamycin resistance and grown overnight in Luria Bertani (LB) broth with 0.1 mg/ml kanamycin.These constructs encode for the first 300 amino acids along with an additional hexa-histidine tag at the N-terminal. Constructs (WT and C231S) were confirmed through DNA sequencing.

### Protein expression and purification

*E. coli* strain BL21 codon plus RIL (DE3) harboring the relevant plasmid was grown overnight at 37°C in LB broth medium supplemented with kanamycin (0.1mg/ml) and chloramphenicol (0.034 mg/ml). 1% of this overnight culture was transferred into fresh medium with kanamycin and chloramphenicol and grown until a cell density equivalent to an OD_600_ of 0.6 was reached. Protein expression was induced with a final concentration of 1mM IPTG for 6 h at 30 °C. Cells were then harvested by centrifugation at 4500xg for 10 min at 4°C, washed with MQ water and frozen at −20°C. Over-expressed proteins were purified using Immobilized Metal Affinity Chromatography (IMAC) based on the affinity of the hexa-histidine tag for Ni^2+^ following cell lysis in lysis buffer (50 mM Tris-HCl, 200 mM NaCl, 1mM PMSF, pH 8.0).All subsequent steps were performed at 4°C. Cell lysis was achieved through sonication done thrice for 5 min with a pulse of 5 sec ‘on’ and 5 sec ‘off’, at 30% amplitude. The lysate was centrifuged at 4500xg for 30 min at 4°C. The supernatant was then applied to Ni^2+^-NTA (Nickel-Nitrilotriacetic acid) sepharose column (GE Healthcare, USA) pre-equilibrated with the lysis buffer at a flow rate of 0.2 ml/min. The protein was eluted in 50 mM Tris-HCl, 200 mM NaCl, 300 mM Imidazole, pH 8.0). The fractions containing the eluted protein were pooled and immediately subjected to buffer exchange using HiPrep™ 26/10 desalting column on an AKTA FPLC purification system (GE Healthcare, USA) into storage buffer (50 mM Tris-HCl, 200 mM NaCl, pH 8.0). The protein was concentrated using Amicon Ultra-15 centrifugal filter units (Millipore, Germany) to around 8 mg/ml. Size-exclusion chromatography was performed at 4 °C using a HiLoad™ Superdex 200pg 16/600 GE preparative column (GE Healthcare, USA). The column was equilibrated with 50 mM Tris-HCl, 200 mM NaCl, pH 8.0. 250 μg protein was loaded into the column and eluted at a rate of 0.5 ml/min. The chromatograms were calibrated with the absorption of the following proteins at 280nm: Ferritin (440kDa), Aldolase (158kDa), Conalbumin (75kDa), ovalbumin (44kDa) and Ribonuclease A (13.7kDa). Over-expression and purification of PTP-PEST was confirmed through SDS-PAGE. Protein concentration estimations were done through Bradford’s assay with bovine serum albumin serving as standard.

### Statistical analysis

All experimental data are represented as mean ± SEM for a minimum of three independent experiments. Statistical evaluation was performed using Student’s *t*-test or one-way ANOVA, followed by Tukey’s multiple comparison post hoc test, using GraphPad Prism version 6.0 software for Windows (GraphPad Prism Software Inc. San Diego, CA, USA). *p* value < 0.05 was considered to be statistically significant.

## Results

### Hypoxia enhances PTP-PEST protein levels and enzyme activity

To determine the effect of hypoxia (1% oxygen) on protein levels and enzyme activity of PTP-PEST in endothelial cells, HUVECs were cultured as monolayer and exposed to hypoxia for different time points. As can been seen in figure 1A and summarized in figure 1B, immunoblotting for PTP-PEST demonstrated a significant increase in protein levels from 3 hour onwards and it was sustained till 24 hours. Increase in protein levels of HIF-1α confirmed induction of hypoxia in these cells. Further, to determine whether this effect of hypoxia on PTP-PEST expression is an endothelial specific phenomenon, or is it also observed in other cell types, we checked for changes in PTP-PEST expression in response to hypoxia in cell lines such as HEK293, HASMC (human aortic smooth muscle cell), HeLa (human cervical epitheloid carcinoma cell line) and Huh7 (hepatocyte derived). As seen in figure 1C, hypoxia promotes PTP-PEST protein expression even in other cell lines, suggesting this to be a universal phenomenon. Equal loading and induction of hypoxia were confirmed for each of the cell lines via immunoblotting for β-actin and HIF-1α respectively (data not shown). We also checked for changes in sub-cellular localization of PTP-PEST if any in response to hypoxia by immunofluorescence imaging. Cytoplasmic localization of PTP-PEST was observed in normoxia which did not alter upon hypoxia treatment (<u>Supplementary figure 1</u>).

**Figure 1.**
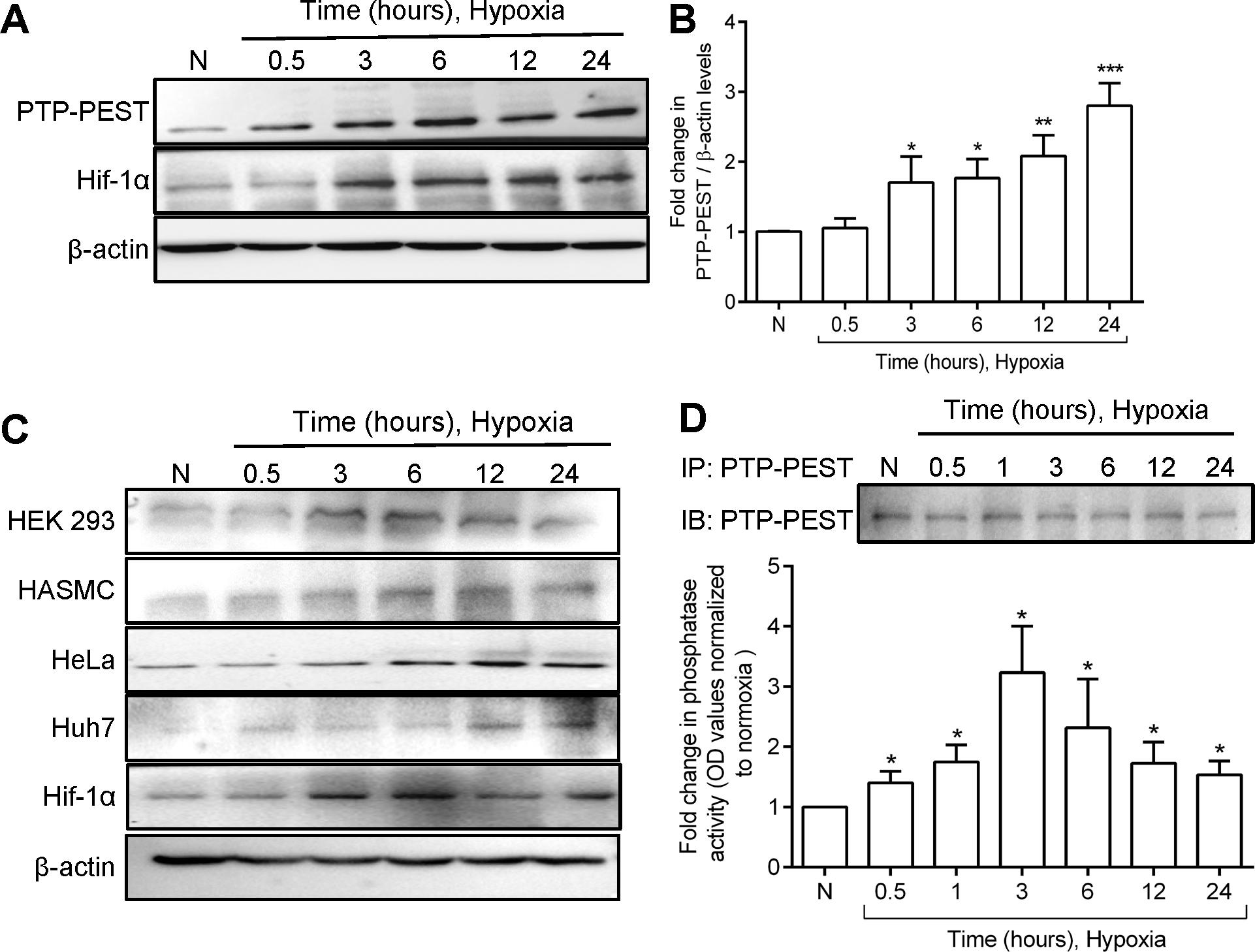
Hypoxia-induced PTP-PEST protein level and enzyme activity. (A) Representative western blot demonstrating effect of hypoxia on protein expression of PTP-PEST and HIF-1α in HUVECs. (B) Bar graph summarizing data for PTP-PEST protein expression for 10 independent experiments. (C) Representative Western blot depicting effect of hypoxia on protein expression of PTP-PEST in HeLa, HASMC HEK293 and Huh7 cell lines. (D) Bar graph summarizing data for PTP-PEST immunophosphatase assay in response to hypoxia (n=3). Inset: representative Western blot panel confirming equal pulldown of immunoprecipitated PTP-PEST. Bar graph represents data as mean ± SEM. (*p<0.05, **p<0.01 and ***p<0.001versus normoxia).

Since endothelial cells by virtue of their location are one of the early responders to changes in oxygen tension and are resilient to hypoxia to promote adaptive angiogenesis [19;20], we employed primary endothelial cells (HUVECs) as model cell system to determine the role of PTP-PEST in hypoxia mediated cellular responses. We set out to determine if hypoxia influences the catalytic activity of PTP-PEST. Equal amount of endogenous PTP-PEST was immuno-precipitated from endothelial lysate and was subjected to immuno-phosphatase assay. As seen in figure 1D, hypoxia induced a significant increase in phosphatase activity of PTP-PEST with the activity being maximal at 3 hours compared to normoxia treatment. Thus, hypoxia increases both protein levels and enzyme activity of PTP-PEST.

### PTP-PEST interacts with AMPKα

To understand the functional relevance of hypoxia mediated enhanced expression and activity of PTP-PEST, it was necessary to recognise its binding partners. Most of the reported binding partners of PTP-PEST are involved in cell adhesion, migration, and cytoskeletal reorganization. We wanted to identify binding partner of PTP-PEST specifically in endothelial cells exposed to hypoxia and normoxia. For this, HUVECs were transduced with N-terminal GST tagged PTP-PEST lentiviral particles. Post-transduction same pool of cells was split and exposed to either normoxia or hypoxia for 24 hours. Equal amount of GST tagged PTP-PEST was pulled down *via* immunoprecipitation using GST antibody and the co-immunoprecipitated proteins were subjected to LC/MS/MS analysis as described in methods section. The protein IDs obtained after mass spectrometry (MS) data were filtered for non-specific binding partners by removal of proteins appearing in MS of the isotype control sample. Further, MS contaminants as per the common repository of adventitious proteins (https://www.thegpm.org/crap/) as well as proteins which are a part of the sepharose bead proteome were excluded from the analysis [21]. Proteins unique only to normoxia or hypoxia were subjected to analysis by gene ontology program PANTHER. The prominent binding partners observed solely in normoxia were signalling molecules, transcription factors and cytoskeletal proteins (figure 2A). Whereas, binding partners such as chaperones, hydrolases and oxidoreductases appeared exclusively in hypoxia (figure 2A). Some of specific binding partners observed in normoxia and hypoxia are listed in figure 2B. In hypoxia, we found proteins like OTUB1, PGK1 and PKC epsilon, which are known to play an important role in regulating autophagy interacting with PTP-PEST. Surprisingly, proteomic studies revealed AMPKα_1_ and α_2,_the catalytic subunits of 5’-AMP-activated protein kinase(AMPK) as interacting partners of PTP-PEST in normoxia (figure 2B). Co-immunostaining of PTP-PEST and AMPKα followed by immunofluorescence imaging demonstrated that both PTP-PEST and AMPKα indeed co-localize in cytoplasm of HUVECs (figure 2C).

**Figure 2.**
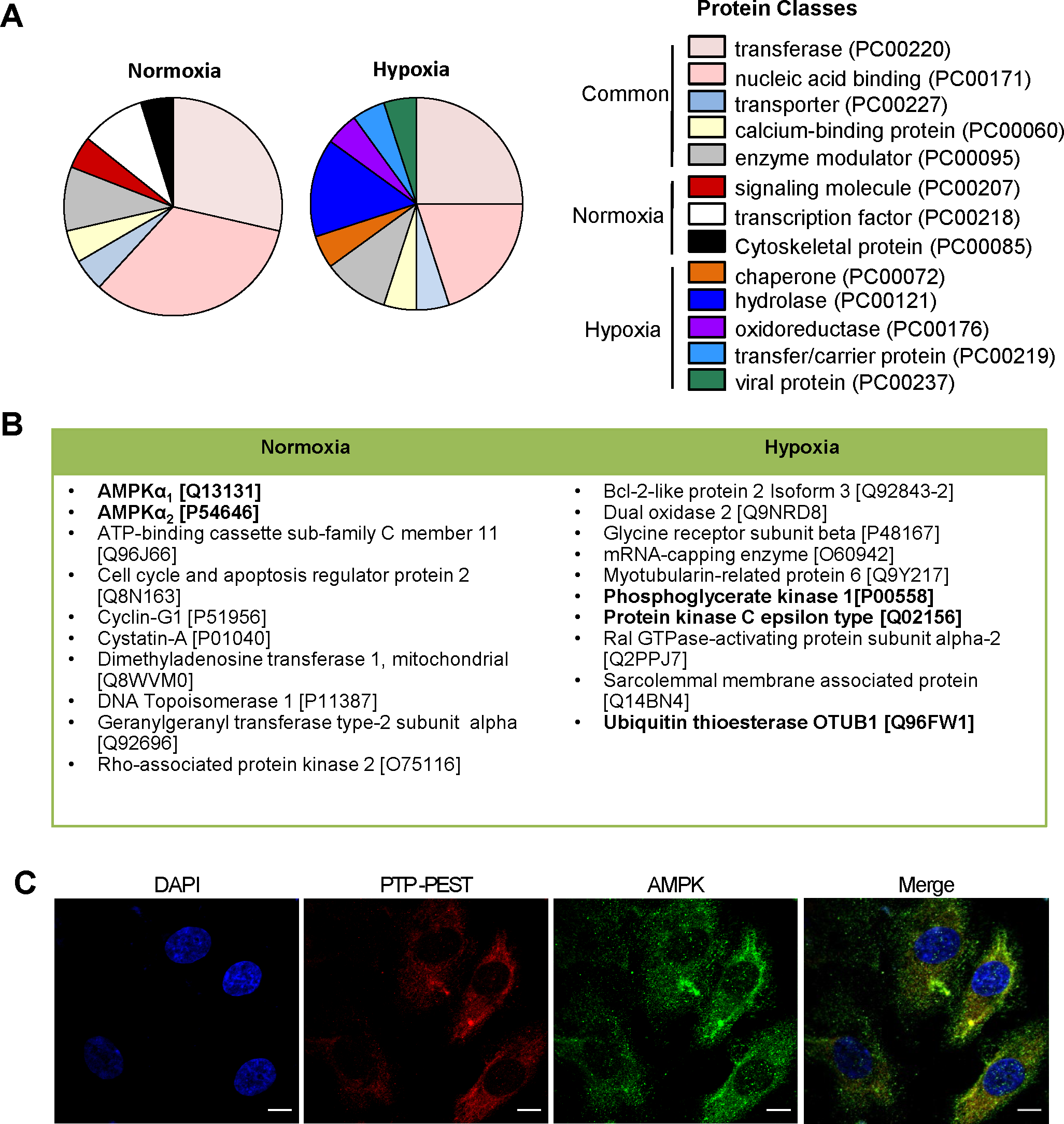
Proteomics analysis of protein IDs obtained from LC/MS-MS data. (A) Pie chart representing classes of interacting protein partners of PTP-PEST in normoxia and hypoxia. (B) List of proteins interacting with PTP-PEST exclusively either in normoxia or in hypoxia. Uniport IDs of binding proteins is listed in parentheses. (C) Representative Immunofluorescence image demonstrating co-localization of PTP-PEST and AMPKα in primary endothelial cells.

Intriguingly, our proteomics data revealed that interaction of PTP-PEST with AMPKα was lost upon hypoxic treatment. In order to validate the interaction of PTP-PEST with AMPKα subunits, we performed co-immunoprecipitation experiments. Equal amounts of endogenous PTP-PEST were pulled down from normoxia and hypoxia exposed HUVECs and immunoblotting was performed for AMPKα. It is worth noting that the AMPKα antibody used for immunoblotting recognizes both the isoforms of α subunit. In concurrence with the proteomics data, we found interaction of PTP-PEST with AMPKα in normoxic condition, which was however abrogated in HUVECs exposed to hypoxia for 24 hours (figure 3A). Equal pull down of PTP-PEST across the two conditions was confirmed by re-probing the blot for PTP-PEST.

**Figure 3.**
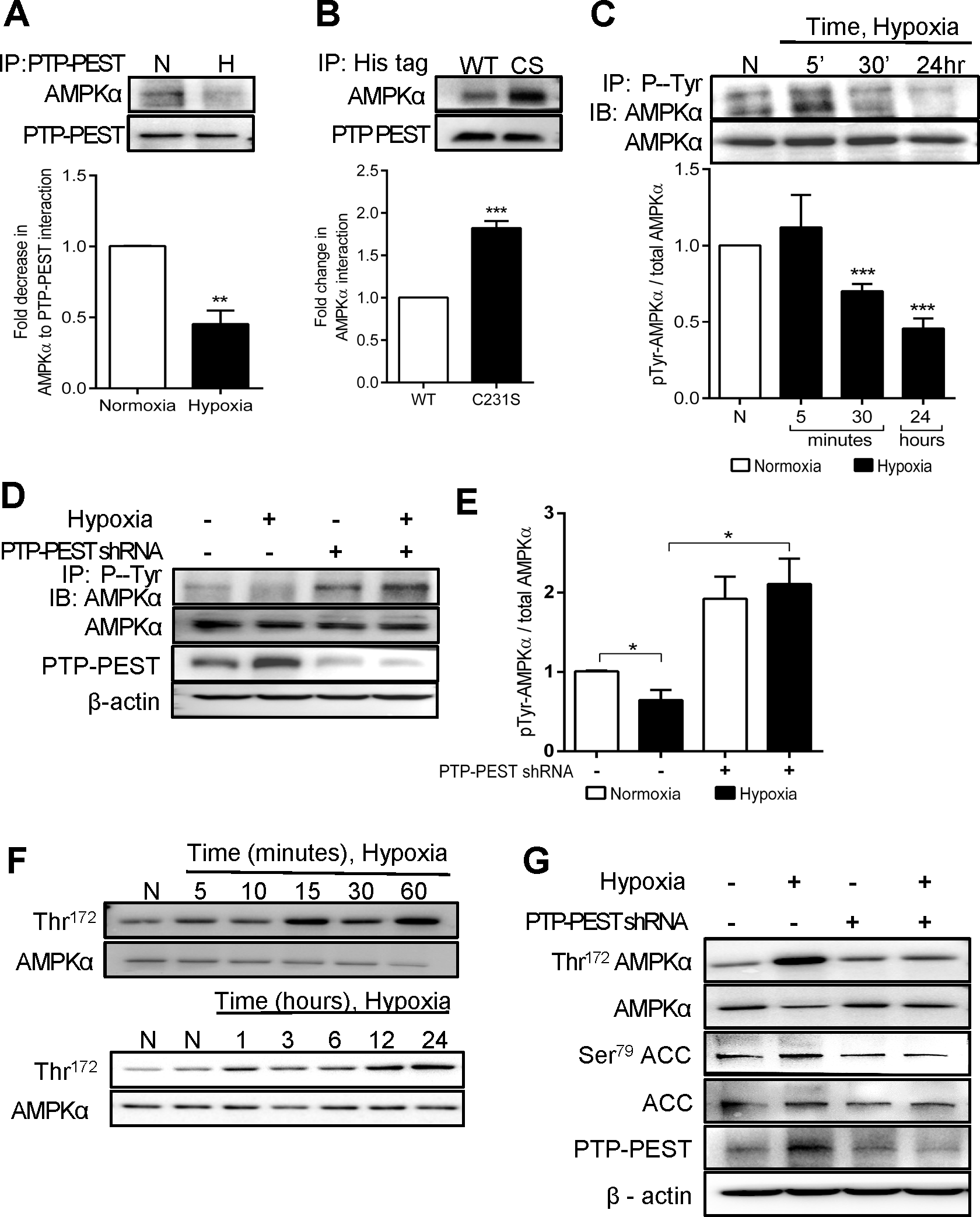
AMPKα is an interacting partner and substrate for PTP-PEST. (A) Representative Western blot and bar graph depicting co-immunoprecipitation of AMPKα with PTP-PEST from HUVECs exposed to hypoxia and normoxia. (B) Representative Western blot and bar graph showing co-immunoprecipitation of AMPKα with purified His-PTP-PEST WT (1-300 amino acids) and His-PTP-PEST C231S mutant (1-300 amino acids). (C) Representative Western blot and bar graph demonstrating effect of hypoxia on AMPKα tyrosine dephosphorylation. (D) Representative Western blot demonstrating effect of PTP-PEST knockdown on hypoxia-induced AMPKα dephosphorylation. (E) Bar graph summarizing effect of PTP-PEST knock-down on AMPK dephosphorylation. (F) Representative Western blot depicting effect of hypoxia on AMPKα activation (Thr^172^phosphorylation). (G) Effect of PTP-PEST knockdown on hypoxia-induced AMPKα Thr^172^ phosphorylation and ACC Ser^79^ phosphorylation. In bar graph data is represented as mean ± SEM. (*p<0.05, **p<0.01 and ***p<0.001versus corresponding normoxia).

PTP-PEST consists of an N-terminal catalytic domain and a long C-terminal non catalytic domain, rich in PEST sequences. The C-terminal domain plays a significant role in regulating the enzyme activity of PTP-PEST by facilitating its interaction with either substrates and/or other adaptor proteins. Next, we checked whether the PEST domain is also essential for the interaction of PTP-PEST with AMPKα. We cloned the His tagged wild type (WT) and catalytic inactive (C231S) N-terminal catalytic domain (1-300 amino acids) lacking the C-terminal PEST sequences of human PTP-PEST into pET28a(+) bacterial expression vector. The two proteins were over expressed and purified from *E. coli* BL21 codon plus RIL (DE3) strain (<u>Supplementary figure 2</u>A-C). 100µg of purified WT and C231S mutant proteins lacking the PEST motifs were independently incubated with 500 µg of HUVEC lysate for 8 hours at 4°C. This was followed by immunoprecipitation of PTP-PEST using a His-tag antibody and immunoblotting for AMPKα. Equal pull down of PTP-PEST was confirmed by re-probing the blot for PTP-PEST. We found co-immunoprecipitation of AMPKα with N-terminal catalytic domain of PTP-PEST (figure 3B), which suggests that the interaction of AMPKα with PTP-PEST is not dependent on the C-terminal PEST domain. Moreover, co-immunoprecipitation of AMPKα with the C231S (catalytically inactive) mutant was greater in comparison to WT (figure 3B), suggesting that AMPKα is a likely substrate of PTP-PEST.

### PTP-PEST mediates hypoxia-induced AMPK activation

AMPK is a hetero-trimeric stress responsive serine-threonine kinase known to play an important role in maintaining cellular energy homeostasis as well as autophagy [22]. Next, we wanted to understand the relevance of interaction of AMPKα subunits with PTP-PEST, a tyrosine phosphatase. Since, PTP-PEST interacts with AMPKα *via* its catalytic domain; we were interested in examining whether PTP-PEST can dephosphorylate AMPKα. First, we determined the effect of hypoxia on total tyrosine phosphorylation of AMPKα. We initially tried immunoprecipitating endogenous AMPKα but faced some difficulties. Hence, we resorted to an alternative approach that was employed by Yamada et al [23]. For this, HUVECs were exposed to hypoxia for different time periods. Equal amount of HUVEC lysate following experimental treatment was subjected to immunoprecipitation with Phospho-Tyr-102 antibody, followed by immunoblotting for AMPKα. Tyrosine phosphorylation of AMPKα in HUVECs exposed to hypoxia for 24 hours was lower in comparison to that exposed to normoxia (figure 3C), indicating its dephosphorylation. It is worth noting that the total AMPKα levels in endothelial cells did not change across treatment conditions (figure 3C). Next, we knocked down PTP-PEST using shRNA lentivirus to check its effect on AMPKα tyrosine dephosphorylation in response to hypoxia (24 hr). Scrambled shRNA lentivirus was used as negative control. We found that hypoxia induced AMPKα dephosphorylation was indeed abrogated upon PTP-PEST knock down (figure 3D and E). In fact, in the absence of PTP-PEST, the basal tyrosine phosphorylation of AMPKα itself was increased, while the total protein levels of AMPKα remained unchanged upon PTP-PEST knock down. Thus, PTP-PEST does regulate tyrosine dephosphorylation of AMPKα in response to hypoxia without influencing its protein levels.

AMPK can be regulated both allosterically and by means of post-translational modifications. Phosphorylation at Thr^172^ residing in the kinase domain of α-subunit is an efficient mechanism of AMPK activation [22]. Therefore, next, the effect of hypoxia on AMPK activation in endothelial cells was examined. HUVECs were exposed to hypoxia for different time points and immunoblotting was performed for phosphorylation of Thr^172^. Thr^172^ phosphorylation was seen in response to hypoxia as early as 15 minutes and it was sustained up to 24 hours (figure 3F and <u>Supplementary figure 2</u>D). Since we observed that PTP-PEST mediates hypoxia induced tyrosine dephosphorylation of AMPK, we set out to determine if absence of PTP-PEST also influences Thr^172^ phosphorylation of AMPK. For this purpose HUVECs were transduced with PTP-PEST shRNA and scrambled shRNA lentivirus followed by hypoxia treatment and immunoblotting for phospho Thr^172^ AMPKα. Interestingly, we found hypoxia induced AMPK activation was attenuated in HUVECs upon PTP-PEST knock down (figure 3G). Simultaneously, we also checked for phosphorylation of ACC at Ser^79^ (a known substrate of AMPK) as a read out of AMPK activity. We found that hypoxia-induced phosphorylation of ACC at Ser^79^ was also attenuated in PTP-PEST knock down cells (figure 3G). These observations, in conjunction with loss of interaction between PTP-PEST and AMPKα under hypoxia, demonstrate that AMPKα is a substrate of PTP-PEST, wherein PTP-PEST mediates tyrosine dephosphorylation of AMPKα and regulates its catalytic activity.

### PTP-PEST regulates hypoxia-induced autophagy via AMPK

AMPK is known to regulate autophagy *via* dual mechanism, involving inactivation of mTORC1 and direct phosphorylation of ULK1 at Ser^317^ [24;25]. Since, we observed that PTP-PEST dependent tyrosine dephosphorylation of AMPKα is associated with its activation in response to hypoxia; we next wanted to understand the role of PTP-PEST in hypoxia-induced endothelial cell autophagy. For this we first tested the effect of hypoxia on endothelial autophagy followed by knock down experiments. HUVECs exposed to hypoxia for different time intervals displayed an increase in LC3 degradation (LC3II form) following hypoxia (<u>supplementary figure 3</u>A&B). An increase in LC3II form indicates induction of autophagy since it plays an indispensable role in autophagosome biogenesis. Along with an increase in LC3II levels, other autophagic markers like beclin-1 and phosphorylation of Ser^317^ ULK1 were also enhanced in response to hypoxia (<u>supplementary figure 3</u>A, C and D). In addition, a decrease in Ser^757^ ULK1 phosphorylation, another indicator of autophagy, was also observed (<u>Supplementary figure 3</u>A and E). It should be noted that Ser^757^ dephosphorylation of ULK1 reflects inactivation of mTORC1. We also examined LC3 puncta formation in response to hypoxia by LC3 immunostaining, followed by confocal imaging. As seen in <u>supplementary figure 3</u>F, high accumulation of LC3 puncta representing autophagosomes was observed in HUVECs treated with hypoxia. This led to a significant increase in the number of puncta per cell as well as percent of LC3 positive cells (<u>supplementary figure 3</u>G and H).Treatment with bafilomycin A1 (100nM) further increased the number of LC3 puncta per cell, indicating that autophagy was in progress during hypoxia. Bafilomycin A1 treatment was given for the last 6 hours of hypoxia treatment.

Next, we wanted to determine whether hypoxia-induced autophagy was dependent on PTP-PEST mediated AMPK activation. As seen in figure 4 A and B, increase in hypoxia induced LC3 degradation was significantly attenuated upon knock down of PTP-PEST. Further, we performed LC3 immunostaining in scrambled and PTP-PEST shRNA treated cells in absence and presence of AMPK activator metformin (500 µM). Metformin treatment was given for 24 hours along with hypoxia. As can be seen from figure 4 C-E, hypoxia-induced increase in number of LC3 puncta per cell as well as percent of cells with puncta-like structures were significantly attenuated upon PTP-PEST knock-down. This attenuation in autophagy due to absence of PTP-PEST was however recovered upon metformin treatment concluding that PTP-PEST is necessary for hypoxia-induced autophagy *via* AMPK activation (figure 4 C-E).

**Figure 4.**
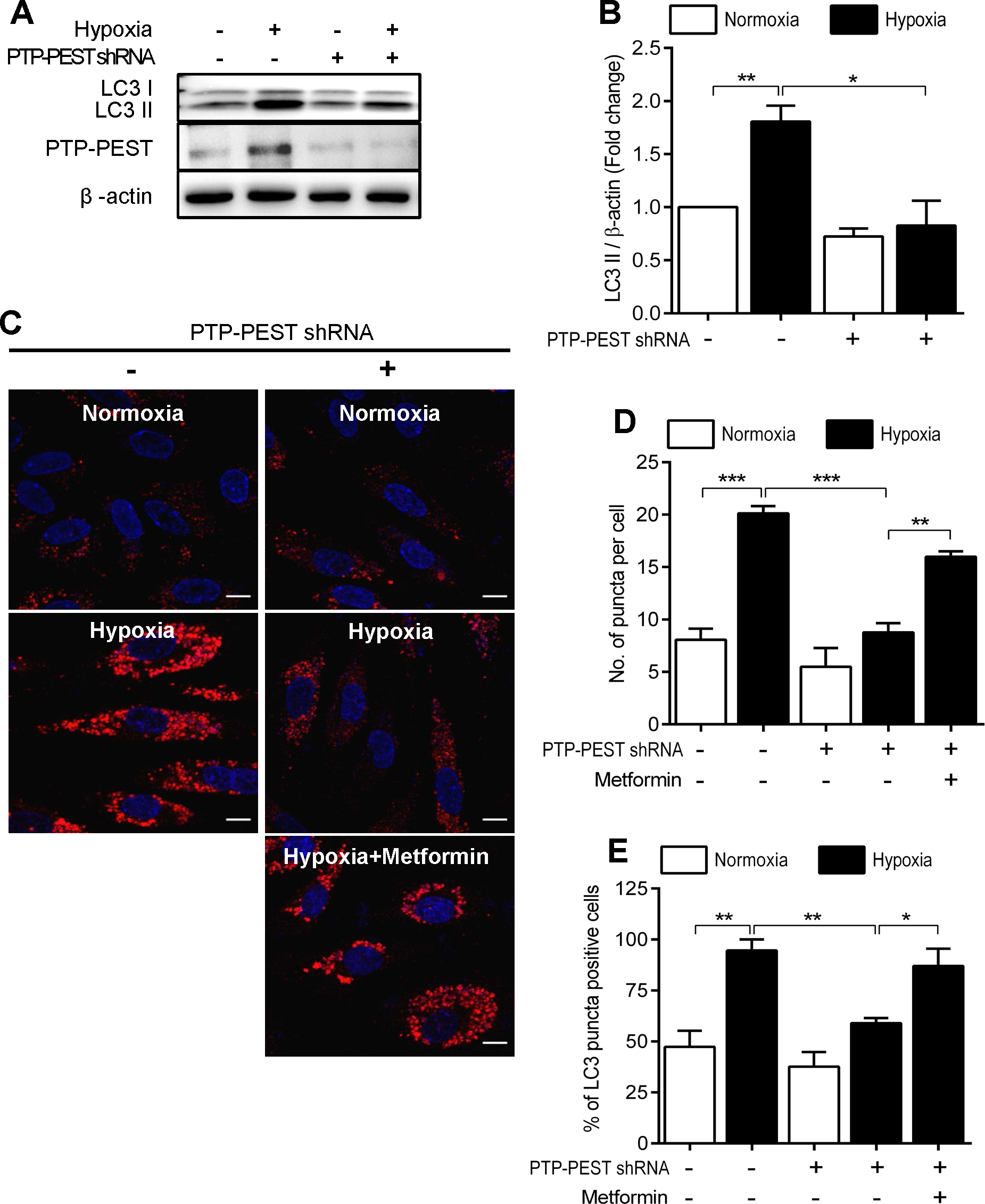
Hypoxia-induced autophagy is dependent on PTP-PEST. (A) Representative Western blotdepicting effect of PTP-PEST knockdown on hypoxia induced LC3 degradation. (B) Bar graph summarizing data of LC3 Western blotting for 5 independent experiments. (C) Representative images showing effect of PTP-PEST knockdown on LC3 puncta formation. (D) Bar graph summarizing data for number of puncta per cell. (E) Bar graph summarizing data for percent of cells with puncta like structures for three independent experiments. In bar graphs data is represented as mean ± S.E.M (*p<0.05, **p<0.01, ***p<0.001 vs corresponding comparisons).

### PTP-PEST regulates hypoxia-induced angiogenesis via autophagy

Hypoxia the principle physiological stimulus for angiogenesis, regulates multiple steps of sprouting angiogenesis including endothelial cell migration, tube formation and vessel branching [26-28]. Additionally, induction of autophagy in response to hypoxia is both a protective survival mechanism as well as an inducer of angiogenesis in endothelial cells [29;30]. We thus examined the effect of PTP-PEST knock-down on hypoxia-induced angiogenic responses such as migration and capillary tube formation. As seen in figure 5A and B, both basal and hypoxia induced endothelial cell migration was attenuated in PTP-PEST knockdown cells, in wound healing experiments. Scrambled shRNA lentivirus was used as a mock transduction. The knockdown of endogenous PTP-PEST was confirmed through Western blotting (figure 5C).

**Figure 5.**
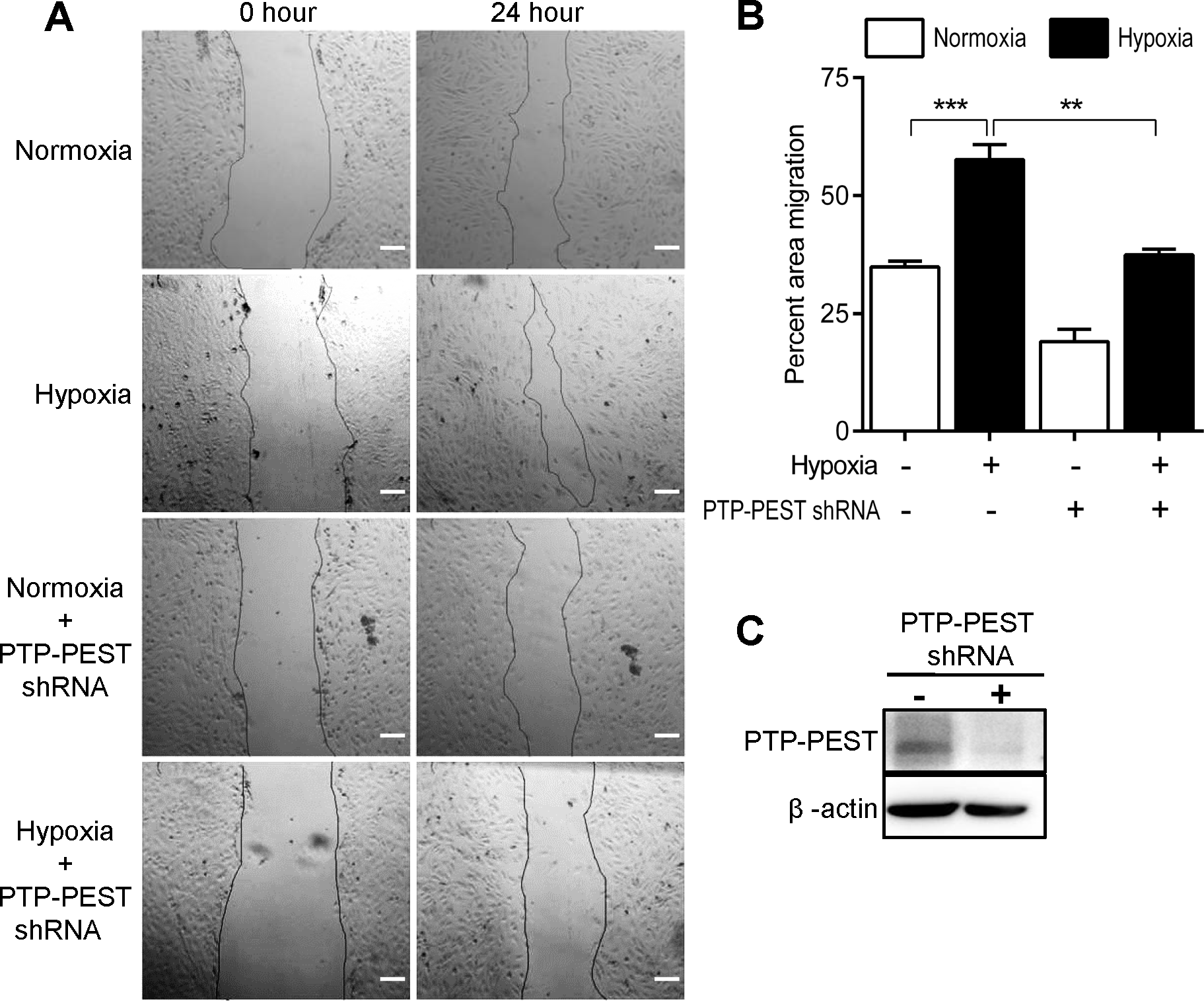
Hypoxia-induced endothelial cell migration is PTP-PEST dependent. (A) Representative images of scratch wound assay demonstrating wound closure by migrating endothelial cells. (B) Bar graph summarizing data as mean ± S.E.M for four independent experiments. (C) Representative Western blot confirming knockdown of PTP-PEST. (**p<0.01 and ***p<0.001 vs corresponding normoxia treatment).

Further, to examine the role of PTP-PEST in hypoxia-induced tube formation *in vitro*, HUVECs were placed on growth factor free Matrigel after transduction with scrambled shRNA or PTP-PEST shRNA lentivirus, and tube formation was assessed under normoxic and hypoxic conditions. Tubular network was quantified after 16 hours of incubation *via* Image J software. We observed an increase in the number of segments, number of junctions as well as in total length of tubes in response to hypoxia (figure 6). Among these effects of hypoxia, PTP-PEST knock-down significantly attenuated number of segments and junctions. PTP-PEST knock down was confirmed through immunoblotting (figure 6E). Interestingly use of rapamycin (200nM), an mTOR inhibitor and autophagy inducer, reversed the effect of loss of PTP-PEST by increasing the number of junctions and segments of tubes for PTP-PEST knockdown cells (figure. 6 A-D). Together, these observations indicate that PTP-PEST promotes hypoxia-induced angiogenesis through AMPK dependent autophagy pathway.

**Figure 6.**
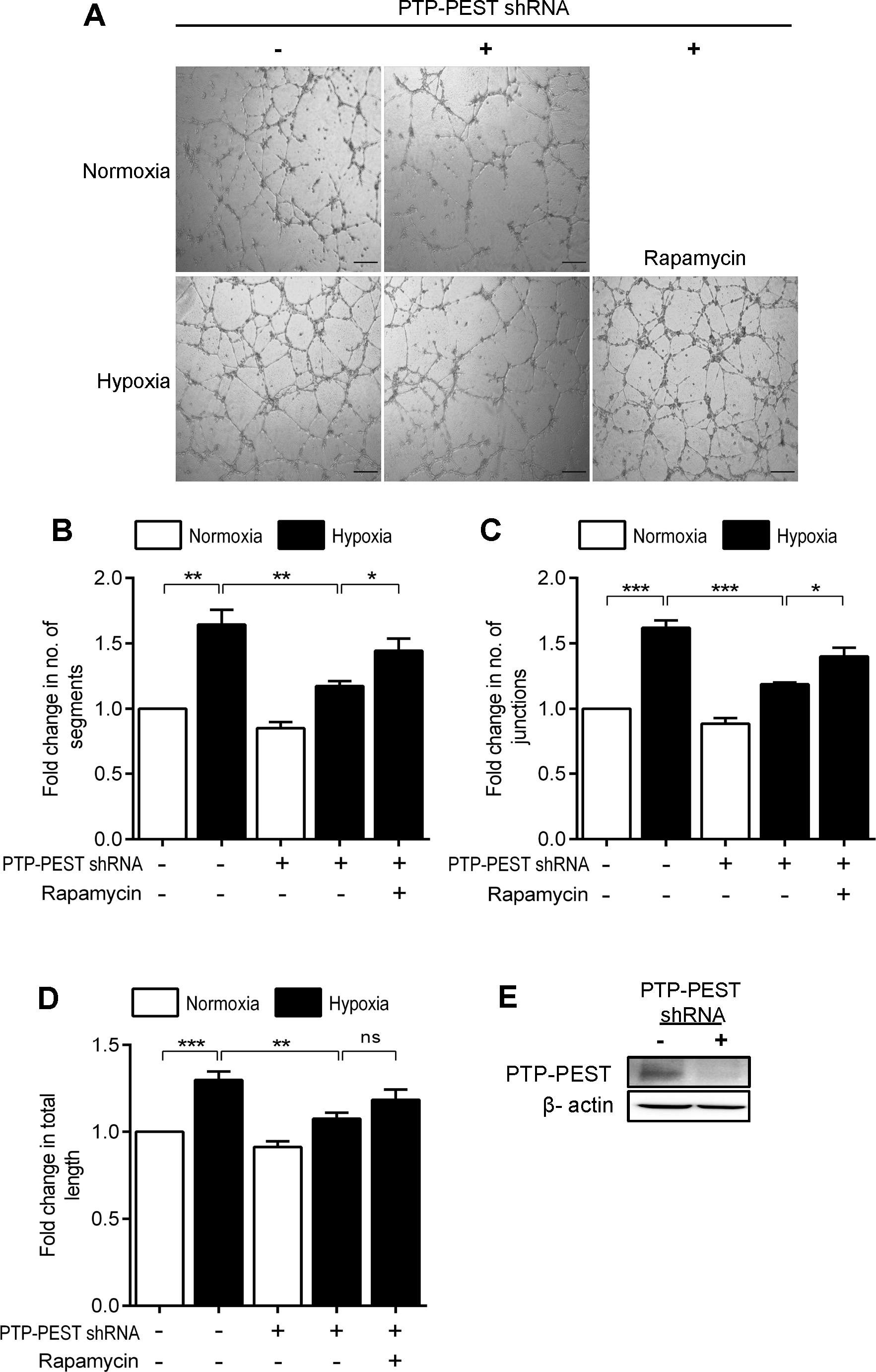
Hypoxia-induced tube formation is PTP-PEST dependent. (A) Representative images for tube formation assay. (B-D) Bar graphs summarizing data for number of segments/tubes, number of junctions and length of tubes. (E) Representative Western blot confirming PTP-PEST knockdown in HUVECs. In bar graphs data is represented as mean ± S.E.M for four independent experiments. (*p<0.05 and **p<0.01 vs corresponding normoxia).

## Discussion

The results of the present study demonstrate that hypoxia increases the expression and catalytic activity of cytosolic tyrosine phosphatase PTP-PEST to promote endothelial autophagy and consequent angiogenesis. These functional effects of hypoxia are dependent on PTP-PEST mediated tyrosine dephosphorylation and activation of 5’-adenosine monophosphate (AMP)-activated protein kinase (AMPK).

As the name suggests, AMPK is a physiological energy sensor that is activated in response to increased intracellular concentrations of AMP under stress conditions of hypoxia, calorie restriction, or exercise. Upon activation, it inhibits energy utilizing anabolic processes and instead, promotes catabolic processes such as fatty acid oxidation, glycolysis and autophagy [22;31]. AMPK holoenzyme is a heterotrimeric complex composed of a catalytic α subunit in complex with two regulatory subunits, β and γ [32]. The kinase domain (KD) of α subunit harbors Thr^172^ residue in the catalytic cleft formed by its N- and C-terminal lobes (figure 7) [33;34]. In an inactive state, the back end of these lobes opposite to the catalytic cleft are held by the evolutionarily conserved Auto-Inhibitory Domain (AID). The AID in turn is connected to a flexible regulatory motif termed as the α-regulatory subunit interacting motif (α-RIM). Multiple crystallography studies have demonstrated that binding of AMP molecules to the γ subunit transmits a conformational change to the α-RIM, allowing it to pull the AID away from the kinase domain to relieve inhibition (figure 7) [35;36]. As a consequence of this long range allosteric regulation, Thr^172^ at the catalytic site is now accessible for phosphorylation by LKB1 or CAMKKβ to enhance its enzyme activity several 100-fold [33;37]. Two isoforms of α subunit (α_1_ and α_2_) exhibiting considerable sequence identity and conformational similarity have been reported [38]. It is worth noting that both these isoforms of α subunit interacted with PTP-PEST under normoxic condition, with the interaction being lost in hypoxia. Interestingly both these isoforms of α subunit are expressed in endothelial cells, where they participate in hypoxia-induced angiogenesis[39;40].

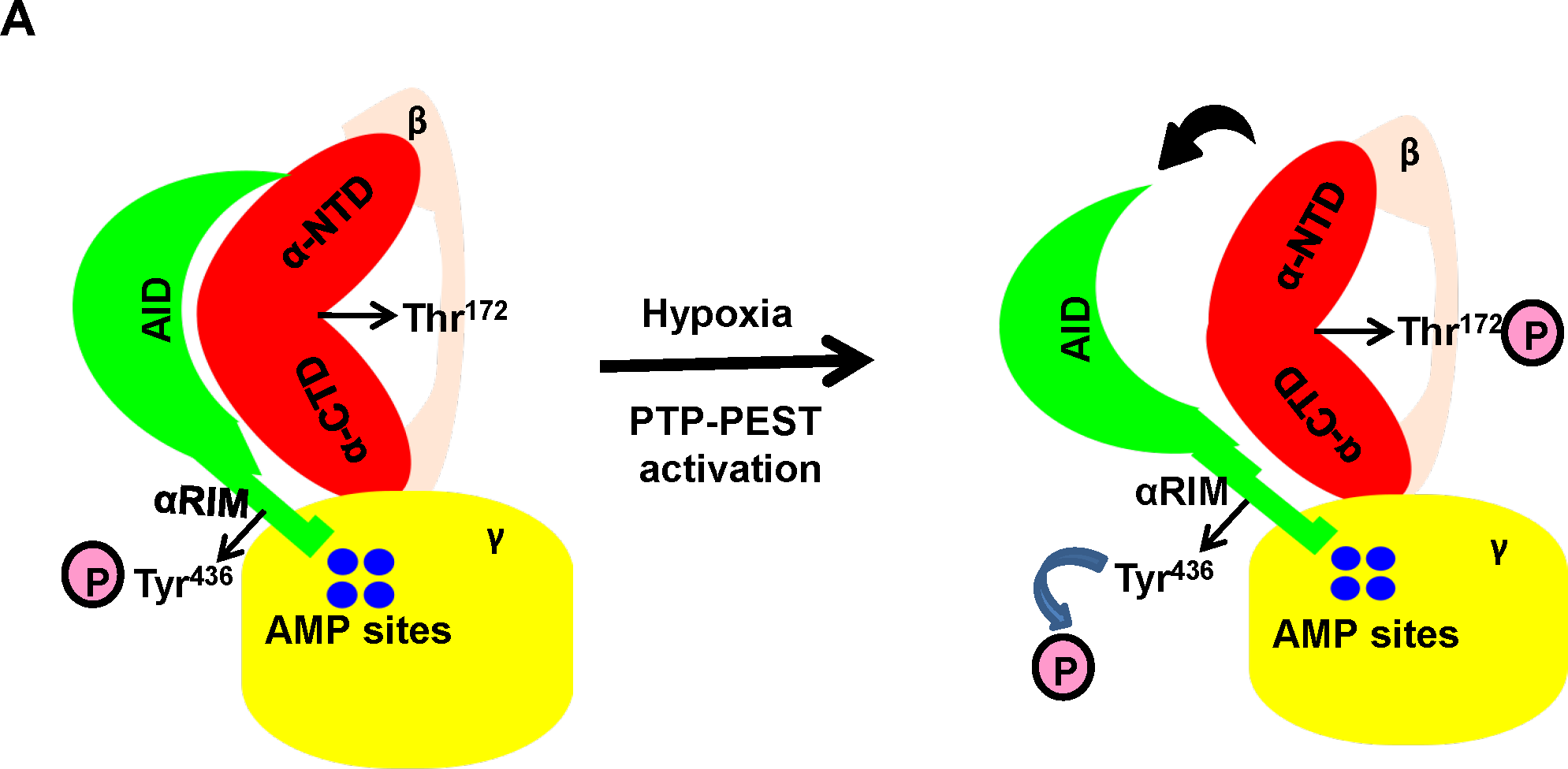

In the current study it was found that hypoxia-induced activation of AMPK was dependent on PTP-PEST. The catalytic domain of PTP-PEST (1-300 aa) was sufficient for interaction with AMPKα, thereby suggesting that AMPKα subunits are PTP-PEST substrates. The endogenous interaction between AMPKα and PTP-PEST was lost in hypoxia. This loss of interaction coincided with tyrosine dephosphorylation of AMPKα. Interestingly, knock-down of PTP-PEST not only enhanced the basal tyrosine phosphorylation of AMPK, but also prevented hypoxia mediated activation of AMPK, as assessed through Thr^172^ phosphorylation. To the best of our knowledge, this is the first study to directly demonstrate regulation of AMPK activity by a cytosolic tyrosine phosphatase. Although not much is known about the regulation of AMPK by tyrosine phosphorylation, a recent study demonstrated that phosphorylation of Tyr^436^ in the α-subunit (figure 7) reduces its catalytic activity by modulating AID-αRIM interaction [23]. It is thus tempting to speculate that PTP-PEST may dephosphorylate this residue of the α-subunit under hypoxic condition to activate AMPK. This possibility however needs to be studied.

One of the major consequences of AMPK activation in endothelial cells is induction of autophagy to promote survival and angiogenesis [19;22;31;41]. This adaptive response imparts resilience to endothelial cells towards stress conditions such as heat shock, hypoxia, shear stress and calorie restriction [42-44]. In fact defective endothelial autophagy is associated with vascular aging, thrombosis, atherosclerosis and even arterial stiffness [42;45]. AMPK activates autophagy either by phosphorylating ULK1 at Ser^317^ or by inhibiting mTORC1 [24;25]. Both these events lead to autophagosome formation. To couple the changes in PTP-PEST and AMPK activity with endothelial cell function, we assessed the consequence of PTP-PEST knock-down on hypoxia-induced autophagy. We found that despite induction of hypoxia, autophagy was attenuated in the absence of PTP-PEST. This effect was however rescued by AMPK activator metformin demonstrating that AMPK activation lies downstream of PTP-PEST in the signalling cascade. Apart from activating AMPK, it was interesting to note that PTP-PEST also interacted with other proteins such as ubiquitin thioesterase (OTUB1), protein kinase C-ξ (PKC-ξ), myotubularin related protein 6 (MTMR6) and sarcolemmal membrane associated protein (SLAMP), all of which are either directly involved in autophagy or its regulation [46-48]. Thus, the current findings also identify hitherto unreported role of PTP-PEST in regulating endothelial autophagy.

PTP-PEST is an efficient enzyme with a K_cat_/K_m_ ≥ 7×10^6^ M^−1^s^−1^ [49;50]. A key finding of the current study is that hypoxia increases both expression and activity of PTP-PEST. It is presently unclear how hypoxia brings about these effects, but a preliminary CONSITE based *in silico* analysis of the putative promoter region of PTP-PEST indicates presence of multiple HIF-1α binding sites (not shown), suggesting a possible increase in PTP-PEST transcription in response to hypoxia. Alternatively, hypoxia may enhance the protein stability of PTP-PEST. A great deal is known about the ability of the ‘PEST’ sequences to regulate protein stability since they are capable of roping in ubiquitin ligases [51]. Post-translational modifications of proline rich ‘PEST’ sequences such as phosphorylation or prolyl hydroxylation assist in these interactions [52;53]. HIF-Prolyl Hydroxylases (PHDs) in presence of oxygen, hydroxylate proline residues to recruit von-Hippel Lindau (vHL) ubiquitin ligases [54]. However, under hypoxic condition they fail to hydroxylate prolines and thus fail to induce protein degradation. In fact this mode of regulation is also responsible for increasing the stability of HIF-1α under hypoxia. The fact that we observed an increased interaction of deubiquitinase OTUB1 with PTP-PEST during hypoxia also supports the notion that hypoxia may increase the stability of PTP-PEST protein.

Increased activity of PTP-PEST could occur as a result of several events, including, changes in sub-cellular localization, protein-protein interactions or even post-translational modifications in response to hypoxia. We did not observe any change in sub-cellular localization of PTP-PEST in response to hypoxia (supplementary figure 1). However, it is possible that PTP-PEST may undergo serine, threonine or tyrosine phosphorylation upon induction of hypoxia. Indeed phosphorylation of Ser^39^ is known to reduce the substrate specificity and activity of PTP-PEST [8]. Likewise, Tyr^64^, an evolutionarily conserved residue in the ‘P-Tyr-loop’ of PTP-PEST is reported to be essential for its catalytic activity [49]. In contrast, Ser^571^ phosphorylation enhances its substrate binding [16]. Whether hypoxia induces serine, threonine or tyrosine phosphorylation of PTP-PEST to modulate either its stability and/or activity is the subject of ongoing investigation in our laboratory.

By virtue of its ability to regulate important cellular processes such as cell adhesion and migration of numerous cell types including embryonic fibroblasts, endothelial cells, dendritic cells, T cells or macrophages, PTP-PEST plays an essential role in cardiovascular development, tissue differentiation and immune function [8;9;13]. Yet the role of PTP-PEST in cancer is controversial with some regarding it as a tumor suppressor while others associate it with metastasis and tumor vasculature [8;15;16]. Our observations of PTP-PEST regulating hypoxia-induced autophagy in native endothelial cells identifies a new functional role of this phosphatase in endothelial physiology. Whether this role of endothelial PTP-PEST is of consequence in tumor angiogenesis under transformed settings needs to be elucidated in future studies. In conclusion, data presented here demonstrates that hypoxia increases expression and activity of cytosolic PTP-PEST, which in turn activates AMPK to promote endothelial autophagy and angiogenesis.

## Acknowledgement

Authors would like to thank Dr. S. Umadevi, Scientist at Translational Research Platform for Veterinary Biologicals, Tamil Nadu Veterinary and Animal Sciences University, Chennai for her technical assistance with confocal microscopy.

## Author Contribution

Shivam Chanel conceived the idea, performed all cell culture experiments, Western blotting, Immunoprecipitation and Immunophosphatase assays, and analysed the relevant data. Amrutha Manikandan analysed the proteomics data. Nikunj Mehta over-expressed and purified WT-PEST and C231S-PEST proteins from *E.coli*. Abel Arul Nathan performed confocal microscopy. Rakesh Kumar Tiwari isolated and cultured endothelial cells. Samar Bhallabha Mohapatra assisted in protein expression and purification experiments. Mahesh Chandran executed in-gel digestion and LC/MS/MS based proteomics. Abdul Jaleel planned and supervised proteomics and performed LC/MS/MS data analysis. Narayanan Manoj supervised biochemical studies with purified proteins and provided intellectual inputs. Madhulika Dixit, procured funding, planned and supervised the cell culture experiments, provided intellectual inputs and wrote the manuscript.

