## Supplementary figure 1 for "Protein tyrosine phosphatase-PEST (PTP-PEST) mediates hypoxia-induced endothelial autophagy and angiogenesis through AMPK activation"

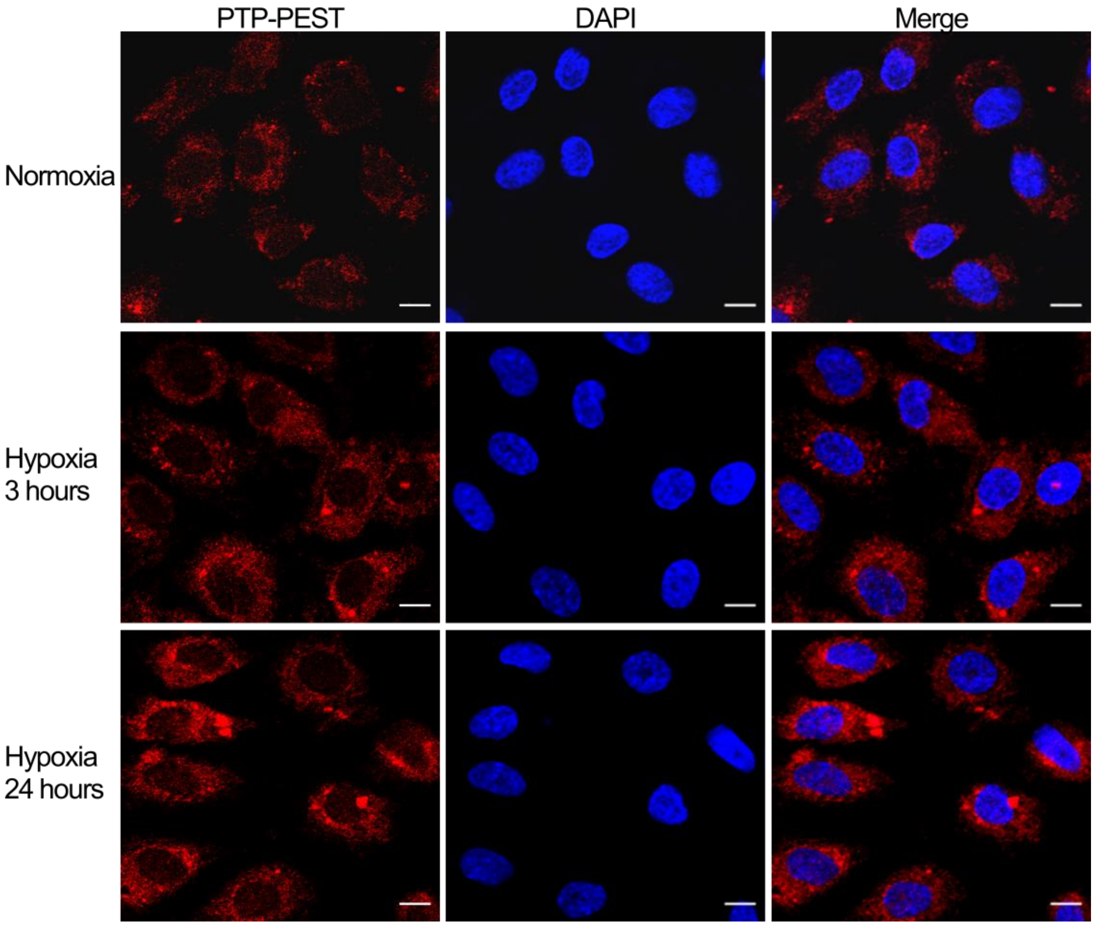


**Supplementary figure 1:** (A) Representative image showing sub-cellular localization of PTP-PEST in response to hypoxia.
