## Supplementary Figure 2 for "Protein tyrosine phosphatase-PEST (PTP-PEST) mediates hypoxia-induced endothelial autophagy and angiogenesis through AMPK activation"

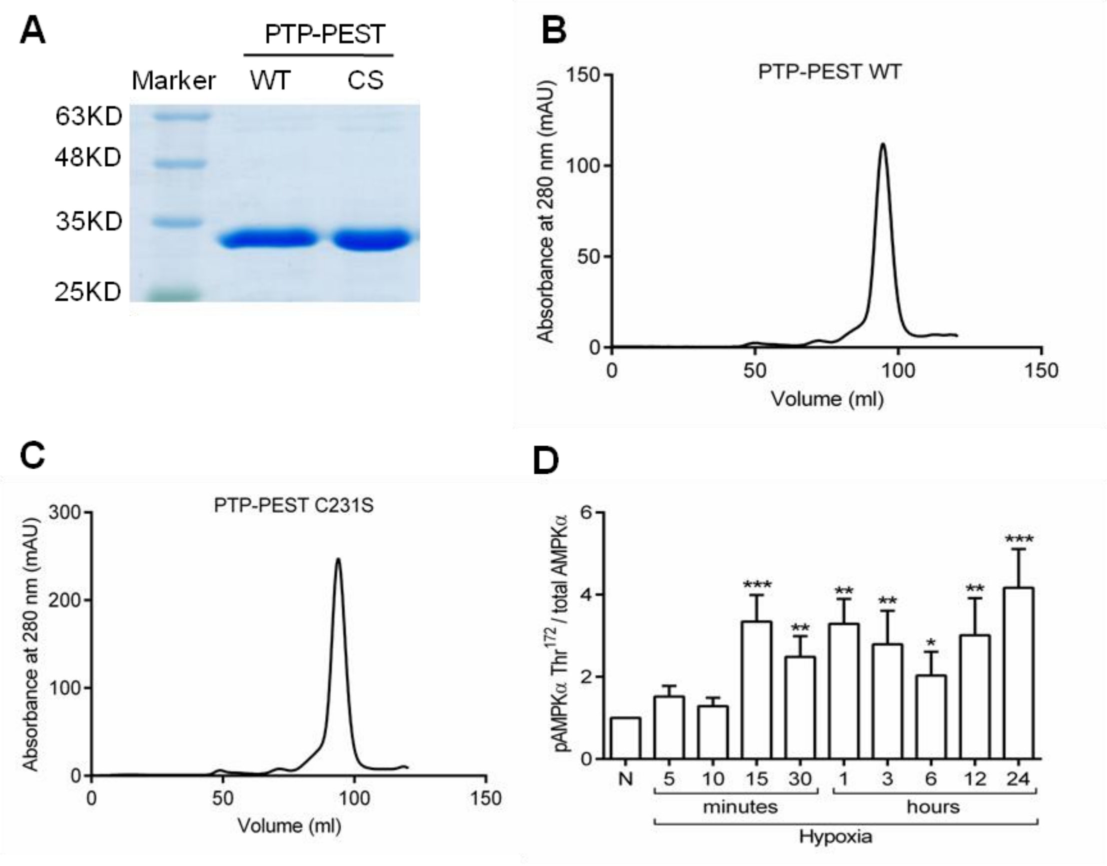


**Supplementary figure 2:** (A) SDS-PAGE of purified His-PTP-PEST WT (1-300 amino acids) and His-PTP-PEST C231S (1-300 amino acids) using $Ni^{2+}$-NTA based IMAC. (B) Size exclusion chromatography of purified His-PTP-PEST WT (1-300 amino acids). (C) Size exclusion chromatography of purified His-PTP-PEST C231S (1-300 amino acids). (D) Bar graph summarizing activation of AMPK in response to hypoxia for five independent experiments as mean ± S.E.M. (*p<0.05, **p<0.01, ***p<0.001 vs corresponding normoxia).
