## Supplementary Figure 3 for "Protein tyrosine phosphatase-PEST (PTP-PEST) mediates hypoxia-induced endothelial autophagy and angiogenesis through AMPK activation"

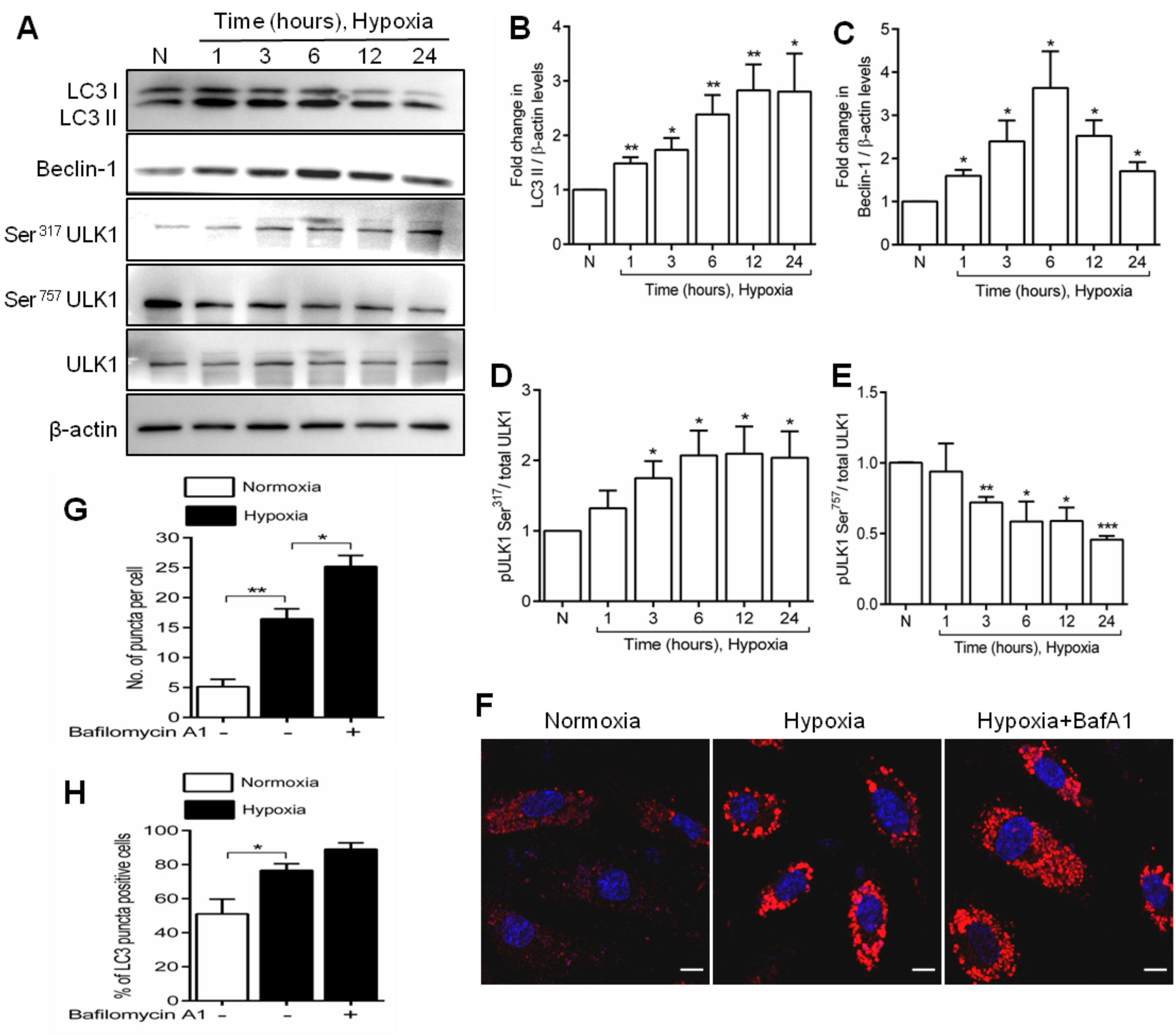


**Supplementary figure 3:**(A) Representative Western blots depicting effect of hypoxia on LC3 degradation, beclin-1 expression, ULK1 $Ser^{317}$ phosphorylation and ULK1 $Ser^{757}$ phosphorylation. (B-E) Bar graphs summarizing data for LC3 degradation, beclin-1 expression, ULK1 $Ser^{317}$ phosphorylation and ULK1 $Ser^{757}$

phosphorylation respectively. (F) Representative images showing effect of hypoxia on LC3 puncta formation in presence or absence of bafilomycin A1. (G) Bar graph summarizing data for number of puncta per cell. (H) Bar graph summarizing data for percent of cells with puncta like structures. In bar graphs data is represented as mean ± S.E.M for a minimum of three independent experiments. (*$p<0.05$, **$p<0.01$, ***$p<0.001$ vs corresponding normoxia).
